# Intrinsic functional connectivity resembles cortical architecture at various levels of isoflurane anesthesia

**DOI:** 10.1101/070524

**Authors:** Felix Fischer, Florian Pieper, Edgar Galindo-Leon, Gerhard Engler, Claus C. Hilgetag, Andreas K. Engel

## Abstract

Cortical activity patterns change in different depths of general anesthesia. Here we investigate the associated network level changes of functional connectivity. We recorded ongoing electrocorticographic (ECoG) activity from the ferret temporo-parieto-occipital cortex under various levels of isoflurane and determined the functional connectivity by computing amplitude envelope correlations. Through hierarchical clustering, we derived typical connectivity patterns corresponding to light, intermediate and deep anesthesia. Generally, amplitude correlation strength increased strongly with depth of anesthesia across all cortical areas and frequency bands. This was accompanied by the emergence of burst-suppression activity in the ECoG signal and a change of the spectrum of the amplitude envelope. Normalizing the functional connectivity patterns showed that the topographical structure remained similar across depths of anesthesia, resembling the functional association of the underlying cortical areas. Thus, while strength and temporal properties of amplitude co-modulation vary depending on the activity of local neural circuits, their network-level interaction pattern is presumably most strongly determined by the underlying structural connectivity.

## Introduction

Ongoing activity of the brain is structured on different spatiotemporal scales. The relationship between spontaneous activity in different brain areas can be quantified through various measures of functional connectivity reflecting different modes of intrinsically generated coupling (Deco et al. 2011; Siegel et al. 2012; Engel et al. 2013). Among these intrinsic coupling modes (ICMs), amplitude envelope correlations (in the following termed envelope ICMs) represent synchronous activity fluctuations of neural populations in two cortical areas at a particular frequency (Engel et al. 2013). Amplitude envelopes are typically modulated at frequencies below 0.1 Hz and show low dependence on behavioral state (Leopold et al. 2003). In the awake state, high envelope ICMs are found between bilateral homologous areas (Nir et al. 2008; Hipp et al. 2012) as well as within functional areas of one hemisphere (He et al. 2008). The spatial specificity of the observed patterns can be increased by excluding effects of volume conduction through orthogonalization (see Methods) (Hipp et al. 2012). Convolution of the amplitude envelope with the hemodynamic response function yields a close estimate of the fMRI blood-oxygen level dependent (BOLD) signal (Liu et al. 2011; Martin 2014). Accordingly, envelope ICMs across two brain areas observed with electrophysiological methods are closely linked to the correlations of their BOLD signals (He et al. 2008; Schölvinck et al. 2010; Keller et al. 2013). Thus, for instance, networks defined by envelope ICMs recorded with MEG show a high similarity to fMRI-resting networks (de Pasquale et al. 2010; Hipp et al. 2012).

Dynamics of ongoing activity are severely altered by general anesthesia, leading to the reversible loss of consciousness. Patterns of cortical activity under anesthesia are not homogeneous, but depend on the concentration of the anesthetic agent. Under light anesthesia, the EEG resembles that of slow-wave sleep (Brown et al. 2010). In deeper anesthesia, brain activity changes to burst suppression, which is characterized by an isoelectric line interrupted by bursts of activity (Llinás and Steriade 2006; Brown et al. 2010). The frequency of bursts decreases with deepening anesthesia, until complete suppression is reached (Brown et al. 2010). Network-level changes caused by anesthetic agents have been examined using various technical and analytical methodologies. Studies focusing on the awake-anesthetized transition have suggested that loss of consciousness is associated with global changes in functional connectivity (Cimenser et al. 2011) and distinct alterations of phase-amplitude-coupling (Lewis et al. 2012; Mukamel et al. 2014). Large-scale envelope ICMs under different anesthesia depths have been studied using fMRI. While some of these studies observe an increase of connectivity with deepening anesthesia (Liu et al. 2013), others contradictorily report a breakdown of coupling (Bettinardi et al. 2015). Micro-scale networks on the cellular level have been characterized using extra- and intracellular microelectrode recordings under different concentrations of anesthetic agents (Llinás and Steriade 2006). Nevertheless, little is known about the corresponding connectivity patterns in mesoscale cortical networks consisting of spatially and functionally related cortical areas.

Here, we characterize envelope ICMs in the visual, auditory and parietal areas of the ferret under different levels of isoflurane anesthesia. We recorded local field potentials from the cortical surface using a custom-built 64-contact electrocorticographic (ECoG) array. Unlike EEG, these recordings are not affected by the distortion of the electrical field in space and frequency by the surrounding tissues (Buzsáki et al. 2012). The anesthetic concentration was varied between 0.4% and 1.6% in steps of 0.2%. For each anesthesia level, envelope ICMs were characterized in different frequency bands using orthogonalized amplitude correlation (Hipp et al. 2012). To account for the different sensitivity of each animal to isoflurane, we applied hierarchical clustering to all connectivity matrices of each frequency band and identified similar coupling modes across animals. The resulting three clusters corresponded to light, medium and deep anesthesia. Overall, the strength of envelope ICMs increased across anesthesia depths. Normalized connectivity patterns, however, remained similar across anesthesia depths, resembling in their topology known patterns of structural connectivity in the cortex.

## Methods

Data presented in this study were collected from six adult female ferrets (*Mustela putorius*). All experiments were approved by the independent Hamburg state authority for animal welfare (BUG-Hamburg) and were performed in accordance with the guidelines of the German Animal Protection Law.

### ECoG-Array

We employed a micro electrocorticographic array co-developed with the University of Freiburg (ECoG-array; IMTEK, Freiburg) covering a large portion of the posterior, parietal and temporal surface of the left ferret brain hemisphere (Rubehn et al. 2009). The probe was designed to record from posterior primary and higher visual areas, primary auditory, somatosensory and parietal areas, located in the medial, suprasylvian and lateral gyrus, respectively (Manger et al. 2002; Bizley et al. 2007). Sixty-four platinum contacts (ø: 250 µm; 0.05 mm2, 7 - 25 kΩ @ 1 kHz) were arranged equidistantly (1.5 mm) in a hexagonal manner on a 10 µm thin polyimide foil. The ∼ 12 x 10.5 mm large foil was subdivided in three ‘fingers’, allowing for a smooth adaptation to the slightly curved surface of the ferret cortex (Fig. 1A).

**Figure 1.**
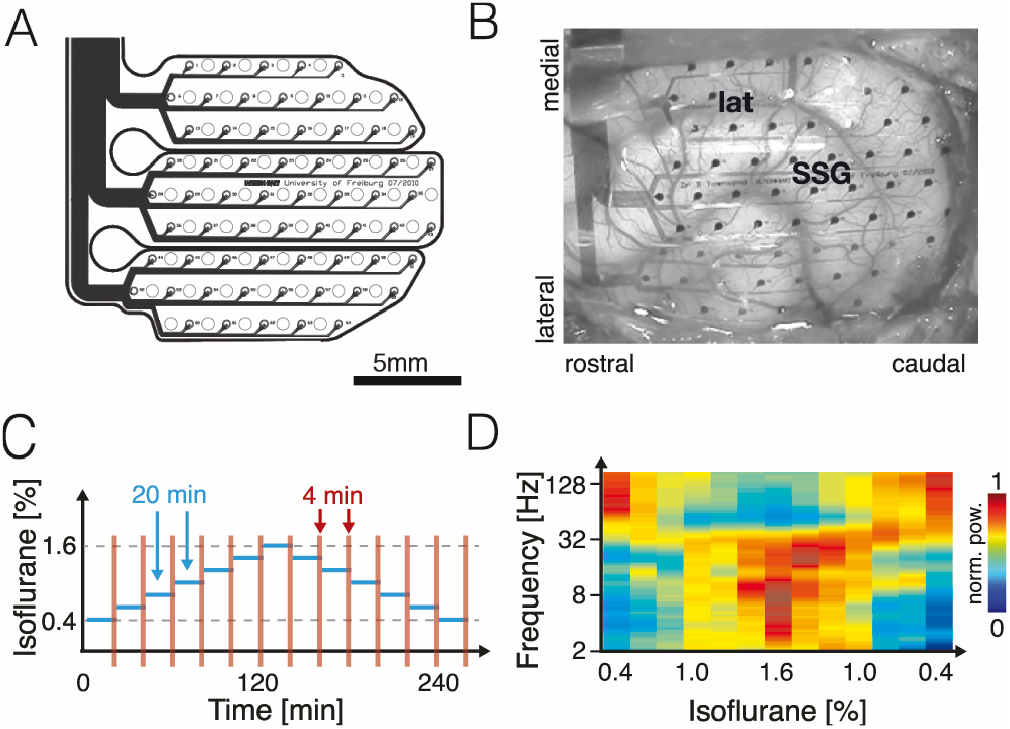
Experimental setting for electrocorticographic recordings in the ferret under different concentrations of isoflurane. (A) A custom-made 64-channel electrode array was used to record the electrocorticogramm. (B) Array in situ. The ECoG-array was designed to cover the temporo-parieto-occipital cortex. (C) During the course of the experiment, isoflurane concentration was varied in 13 steps across seven levels. Each level was kept for 20 minutes, of which the last 4 minutes were chosen for analysis. (D) Average power (normalized to the maximum per frequency) during the course of the experiment for all six animals. While frequencies below around 30 Hz showed a relative increase with higher isoflurane concentrations, higher frequencies displayed the opposite effect.

### Surgery

Animals were initially anesthetized with an injection of ketamine (15 mg/kg) and medetomidine (0.08 mg/kg). After tracheotomy, a Y-shaped glass tube was placed in the trachea to allow artificial ventilation and the administration of isoflurane (0.4-1.6%, 1:2 N_2_O - O_2_ mix; see in more detail below). To prevent dehydration throughout the experiment, a cannula was inserted into the femoral vein to deliver a continuous infusion of 0.9% NaCl, 0.5% NaHCO_3_ and pancuronium bromide (60µg/kg/h). Body temperature was controlled rectally and automatically maintained at 38°C with a homeothermic blanket. After the initial sedation, 1.0% isoflurane was added to the ventilated air to keep anesthesia throughout the following surgical procedures. An anaesthetic monitoring device (Datex-Ohmeda S/5, Datex-Ohmeda Medical Instrumentation Inc., Tewksbury, US) was used to continuously measure inspiratory and expiratory concentrations of isoflurane, MAC, N_2_O, O_2_ and CO_2_ throughout the course of the experiment. In addition, ECG and heart rate was permanently monitored. All data was digitized and logged to disk via the electrophysiological recording systems.

To get access to the left parieto-occipital cortex, first the temporalis muscle was reflected laterally. A saline-cooled and soft-tissue-safe micro-saw (Mectron, Piezosurgery) was used to create a craniotomy (∼13 x 11 mm) over the left posterior cortex. After careful removal of the dura, the ECoG-Array was placed on the cortex. The ECoG-array was covered with artificial dura (Lyoplant/Braun) and the previously extracted bone flap was returned into place and fixed with KLUSTA-KWIK (World Precision Instruments, WPI), a tissue-safe silicon-based two-component glue. Finally, the temporal muscle and the skin were flapped back in anatomical position and provisionally fixed with a suture. This ensured that the cortex would reside in physiological cortico-spinal fluid and avoided cooling of the cortex to nonphysiological temperatures throughout the course of the experiment (Kalmbach and Waters 2012).

After the end of the whole experimental session (∼24h – 36 h), the animal was deeply anesthetized with 5% isoflurane and sacrificed with an intravenous bolus of 3M potassium chloride.

### Data acquisition

The continuous stream of neuronal data (∼4.5 h) was processed with an AlphaLab ‘SnR’ electrophysiological recording system. It was amplified 200-fold, hardware bandpass filtered (0.5 Hz; 2-pole – 8 kHz; 3-pole) and digitized at 44.6 kHz sampling rate. It was then immediately software lowpass filtered (347 Hz), downsampled by factor of 32 (1.4 kHz) and saved to disk. Simultaneously, vital signs and anesthesia parameters as described above were sampled with the same system (2.8 kHz) and also stored to disk.

The data acquisitions for this experiment started between 5 h and 9.5 hours after the initial ketamine anesthesia and after at least 4 hours of isoflurane ventilation. Before the recordings started, the isoflurane level was kept at 0.4 % for at least one hour. During the recording period the isoflurane concentration was altered every 20 minutes in steps of 0.2%, first increasing up to 1.6% and then decreasing again down to 0.4%. This resulted in 13 different conditions of ‘increasing’ and ‘decreasing’ levels of isoflurane. After each dose-change it took about 2-4 minutes until the equilibration of isoflurane concentration between the inspiratory and expiratory airflow was reached.

### Data Analysis

All data analysis was performed with Matlab (The MathWorks Inc.), partially using the fieldtrip-toolbox (Oostenveld et al. 2011) and the chronux-toolbox (http://chronux.org/; Mitra 2008). Offline data were phase preservent bandpass-filtered between 2 Hz and 300 Hz and further downsampled to 700 Hz. A 50 Hz, 100 Hz and 150 Hz notch-filter (1 Hz width, 4-pole butterworth) were applied. Broken contacts, as indicated by extraordinarily low signal amplitudes or uncommon high-amplitude noise, were excluded from further analysis. All data were visually inspected before further analysis to exclude ECG- or other electrical artifacts.

We computed the power spectra with a multitaper (4/1) analysis of all channels and conditions between 2 Hz and 256 Hz at 6 frequency bands per octave. Power in the classical EEG frequency bands – theta (4-8 Hz), alpha (8-13 Hz), beta (13-30 Hz), low gamma (30-64 Hz), high gamma (64-128 Hz) – was calculated as the mean of the corresponding center frequencies.

Envelope ICMs were quantified by computing orthogonalized amplitude correlations. As shown previously, these are not affected by zero-phase lagged components of the channels which might be a result of volume conduction or other, external common noise (for a detailed description of the method, see Hipp et al. 2012). Based on a time-frequency-representation of the two signals, orthogonalization of signal B to A for time t and frequency f was calculated as

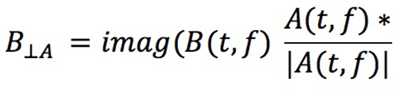

Orthogonalized amplitude correlations where then calculated as

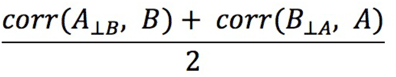

The underlying time-frequency-representation was calculated based on Morlet wavelets. The temporal resolution was set to 1/2 of the maximum frequency’s cycle duration (7.8ms), while the initial frequency resolution was at eight logarithmically spaced bands per octave, resulting in 56 Morlet wavelets of 7 cycles. For further analysis, we calculated the mean correlation coefficients of each channel pair in the theta, alpha, beta, low gamma and high gamma band as specified above.

We analyzed the spectral properties of the amplitude modulation by calculating a multitaper spectrum of the amplitude envelope of each channel. To estimate the amplitude correlation in a given modulation frequency band, we orthogonalized one signal with respect to the other as described above and applied a band-pass filter to both amplitude time courses before calculating the correlation coefficient.

### Assignment of contacts to cortical areas

To account for inter-animal differences in the position of the ECoG-Array and thus make cross-animal comparisons possible, area-level correlation matrices where calculated for cross-animal analysis. Using the sulci as landmarks, a schematic representation of the array position for each experiment was generated relative to a functional map of the ferret cortex (Bizley et al. 2007). We then assigned each contact to a cortical area and calculated the average connectivity values between areas.

### Hierarchical clustering

For each frequency band, we applied agglomerative hierarchical clustering as implemented in Matlab to the area-level coupling matrices of all animals and conditions to identify common network states. To ensure the robustness of the result, we compared the emerging clusters for two distance measures (euclidian and cityblock) and using two linkages (ward and average). An exemplary result is shown in Supplementary Figure 2. We chose to partition the dataset into three clusters. At this level the consistency of the defined clusters across algorithms was high, and the dendrograms showed a clear separation of the clusters from the next level. The results presented here are based on average clustering by Euclidean distance.

**Figure 2.**
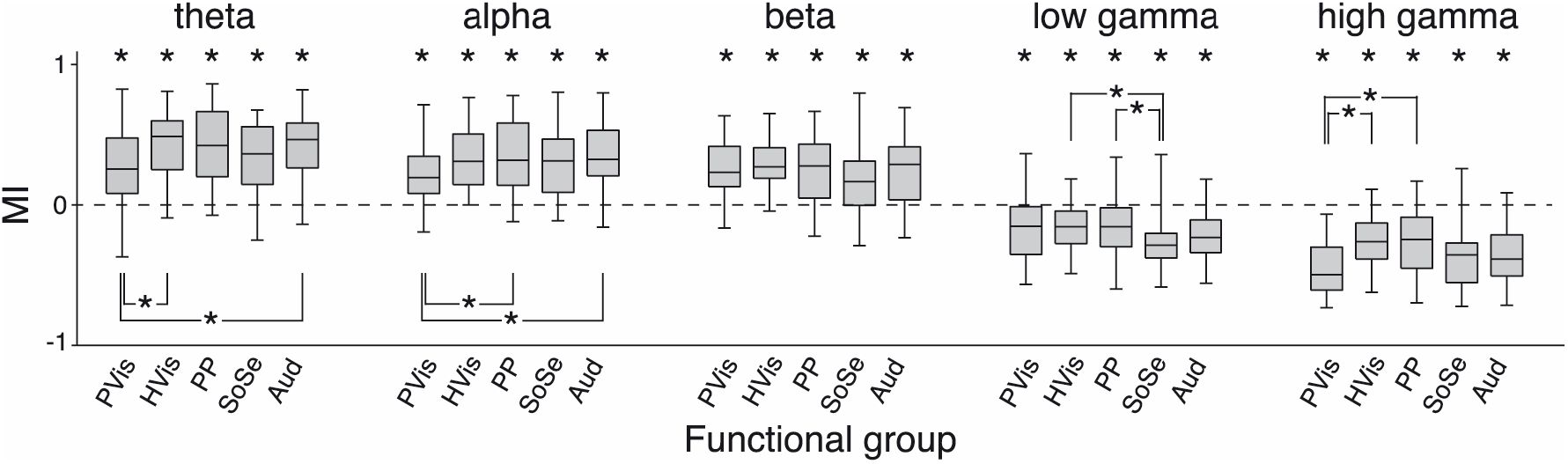
Modulation index (MI) of the power within different groups of cortical areas at 1.6% isoflurane relative to the starting condition at 0.4% isoflurane. In all groups, the power showed significant difference to the starting condition (MI different from 0, ks-test, p<0.05, Bonferroni-Holm-corrected). While power in all groups increased in the theta-, alpha- and beta-band, it decreased in the gamma- and high-gamma band. Abbreviations: PVis - primary visual, HVis - higher visual, PP - posterior parietal, SoSe - somatosensory, Aud - auditory.

### Normalization of correlation values

To examine correlation differences which could not be explained by the distance between contacts, each correlation coefficient in the contact-level connectivity matrix was reexpressed relative to the correlations of all contact pairs with the same inter-contact distance. Contact distances were rounded to the first decimal place. We only considered distances which were represented by at least 25 contact pairs in all animals. All distances above the first one not fulfilling this criterion were excluded. This resulted in 18 distances between 1.5 mm and 9.8 mm (Suppl. Fig. 3). The z-score z = (x-μ)/σ was calculated for each of the correlation values.

## Results

We recorded ongoing cortical activity from the left parieto-temporo-occipital cortex of 6 anesthetized female ferrets using a custom-made 64-channel subdural electrode array (Fig. 1A). Animals where ventilated with a mixture of isoflurane and N_2_O. The isoflurane concentration was varied between 0.4% and 1.6% in steps of 0.2%. Each level was kept constant for 20 minutes to allow the equilibration of a stable network state. The last 4 minutes of these segments were chosen for analysis (Fig. 1C).

### Anesthesia increases low-frequency and decreases high-frequency power

We examined the effect of deepening anesthesia on power change relative to the first data segment in the typical frequency bands. For this purpose, we calculated the modulation index (A-B)/(A+B) within each contact – with ‘A’ being the power of the segment of interest, ‘B’ of the first data segment – scaling the values to a range from -1 to 1. With increasing isoflurane concentration all animals showed a relative increase in theta, alpha and beta power and a relative decrease in gamma power (Fig. 1D).

We next assessed whether there were any topographical differences in the power change occurring from light to deep anesthesia. We pooled the power values across five functional groups (primary visual, higher visual, auditory, posterior parietal and somatosensory) and compared their modulation index at 1.6% isoflurane. For all frequency bands, the power change relative to 0.4% isoflurane was significant for all functional groups (Komolgorov-Smirnov-Test (KS-Test), Bonferroni-Holm-corrected, p<0.05). However, there were only few significant differences between the functional groups (Fig. 2). Overall, the observed relative change in power occurred relatively homogeneously across the whole temporo-parietooccipital cortex. This result is in accordance with other studies showing an occipito-frontal power shift under anesthesia rather than local differences between the areas of the occipital cortex (Cimenser et al. 2011; Supp et al. 2011; Sellers et al. 2013).

### Cluster analysis of envelope ICMs reveals three anesthesia-dependent states

For each 4-minute data segment, we calculated orthogonalized amplitude correlations between all channels. This measure excludes the zero-phase-lag component of the two signals and is thus not susceptible to potential effects of volume conduction (Hipp et al. 2012). Correlation values where calculated on a wavelet-transformed time-frequency-representation and then pooled to the following frequency bands: theta (4-8 Hz), alpha (8-13 Hz), beta (13-30 Hz), low gamma (30-64 Hz), high gamma (64-128 Hz).

For these frequency bands, we then computed coupling matrices reflecting the envelope ICMs at different anesthesia depths. It has long been known from clinical settings that different anesthetic concentrations are needed to elicit the same anesthetic effect in different individuals. We hypothesized that due to inter-individual differences in responsiveness to the anesthetic agent, similar coupling patterns might emerge in the individual animals under different concentrations of isoflurane. Therefore we used hierarchical clustering to identify similar coupling patterns for each frequency band across animals and anesthesia depths.

For each frequency band, we calculated the envelope ICM coupling matrix for each data segment and animal. Amplitude correlation values where averaged within 14 cortical areas covered by the ECoG-probe in all animals. This eliminated inter-individual differences of the contact positions and number of recording sites per cortical area, allowing cross-animal comparison. For each frequency band, the matrices were vectorized and subjected to hierarchical clustering (Suppl. Fig. S1). Based on the dendrograms, we chose to partition the dataset into three clusters corresponding to network states defined by functional connectivity under light, medium and deep anesthesia (Fig. 3A). The deep anesthesia connectivity state was consistent across all frequency bands but appeared in only four of the six animals. In contrast, the light and medium anesthesia connectivity pattern was observed in all animals. They were nearly identical for theta, alpha and beta band. In the low- and high gamma band, the medium anesthesia connectivity state occurred at lower isoflurane concentrations than in the lower frequency bands. This indicates a frequency-band related difference in networklevel effects of isoflurane.

**Figure 3.**
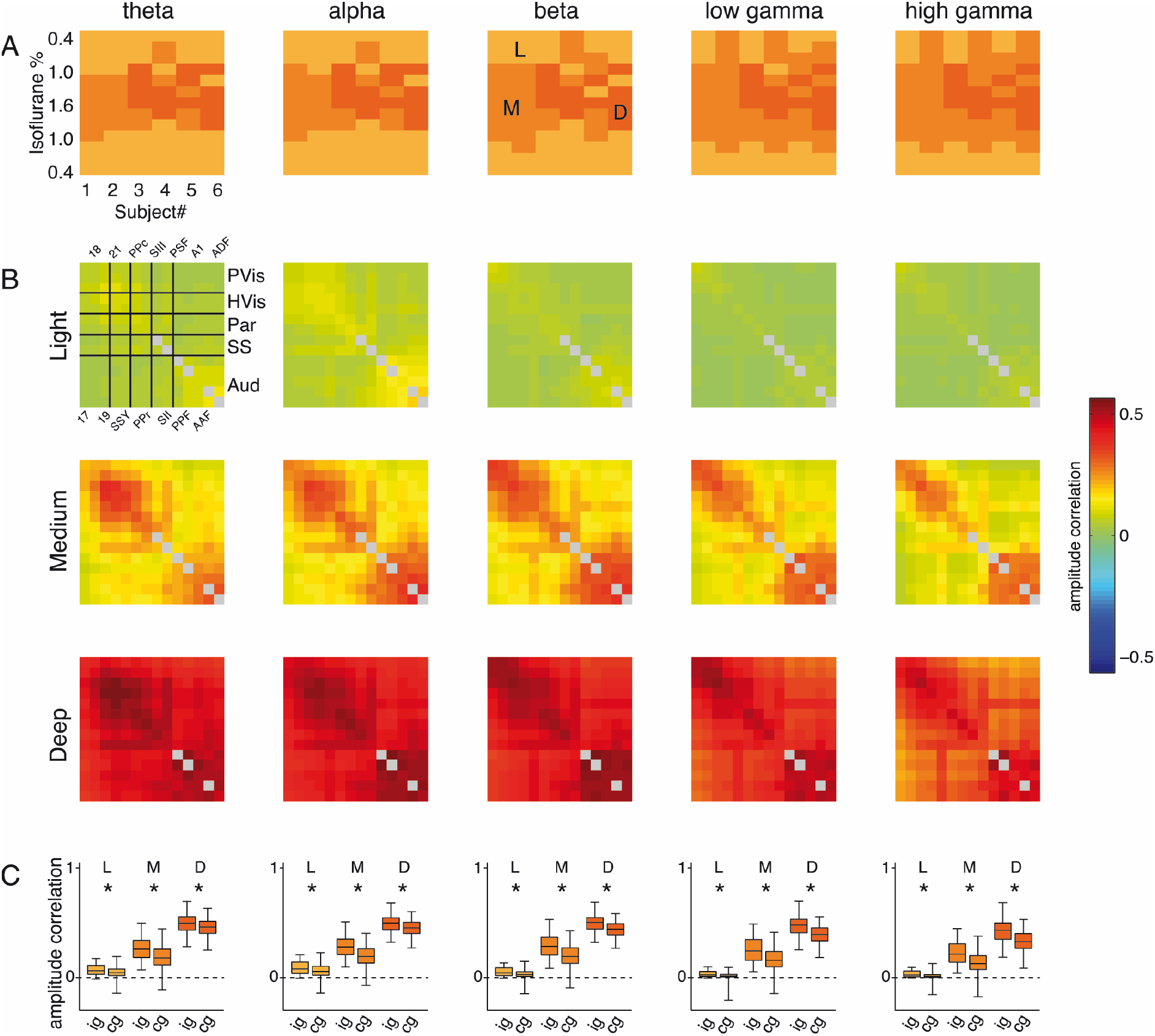
Network states defined by envelope ICMs at different anesthesia depths. (A) Hierarchical clustering of the coupling matrices of individual animals and conditions within each frequency band revealed the emergence of distinct network states under light (L), medium (M) and deep (D) anesthesia. (B) Mean coupling matrices of the three network states. Areas which where only represented by one contact in some of the animals and for which no intra-areal coupling could be calculated, are plotted in grey. (C) Mean correlations within (ig) and across (cg) groups of areas for the three network states. Within-group correlation exceeded cross-group correlation for all frequency bands in all three states (p<0.05, ks-test, Bonferroni-Holm-corrected). Abbreviations: PVis - primary visual; HVis - higher visual; Par - parietal; SS - somatosensory; Aud - auditory; A1 - primary auditory cortex; AAF - anterior auditory field; ADF - anterior dorsal field; PPc - central posterior parietal cortex; PPF - posterior pseudosylvian field; PPr - rostral posterior parietal cortex; PSF - posterior suprasylvian fields; SII - sencondary somatosensory cortex; SIII - tertiary somatosensory cortex; SSY - suprasylvian field.

Inspection of the raw ECoG-signal for the three connectivity states revealed patterns known from single-cell and EEG recordings (Steriade et al. 1994). Example traces are shown in Suppl. Fig. S2. The data segments in the light anesthesia state mostly revealed continuous activity patterns, which in some animals were overlaid by bursts of higher-frequency activity. For the medium anesthesia state frequent bursts with short periods of suppression were observed and for deep anesthesia groups of bursts interrupted by long suppression intervals (Suppl. Fig. 2). At the lowest isoflurane dose of 0.4%, three animals already showed burst suppression patterns. At the highest concentration of 1.6%, all animals had entered burst suppression, four displaying the deepest burst suppression state.

For each anesthesia network state, we calculated the area-level coupling matrices averaged across animals (Fig. 3B). Within each frequency band, global correlation strength increased from the light to the deep anesthesia network state for all fields of the coupling matrix. A KSTest between the clusters confirmed that this difference was significant for all connections in the matrix (Bonferroni-Holm-corrected, p<0.05).

The structure of the coupling matrices suggested that particularly high correlations occurred among functionally closely related areas. To test this hypothesis, we pooled the single-contact correlation values to five functional groups (primary visual, higher visual, posterior parietal, auditory, and somatosensory) and compared correlation values within these functional groups to those across functional groups (Fig. 3C). Within-functional group correlations were consistently higher than across functional group correlations for all network states and frequency bands (KS-Test, Bonferroni-Holm-corrected, p<0.05).

### Overall strength of envelope ICMs shows dependence on distance and frequency range

To capture general properties of envelope ICMs within the three network states, we examined their dependence on frequency and spatial distance of the contacts. For this purpose, we calculated the mean correlation values across all electrode combinations and within six contact distance categories for the three clusters (Fig. 4). For all three anesthesia-dependent network states, correlation showed a drop-off as a function of distance and frequency. In the light anesthesia network state, envelope ICMs showed a peak in the alpha-frequency range and continuously diminished with increasing frequency. During medium depth anesthesia, the highest correlation values where observed in the beta band; during deep anesthesia, mean envelope ICM was similar in theta, alpha and beta band, and only dropped within the gamma and high gamma band.

**Figure 4.**
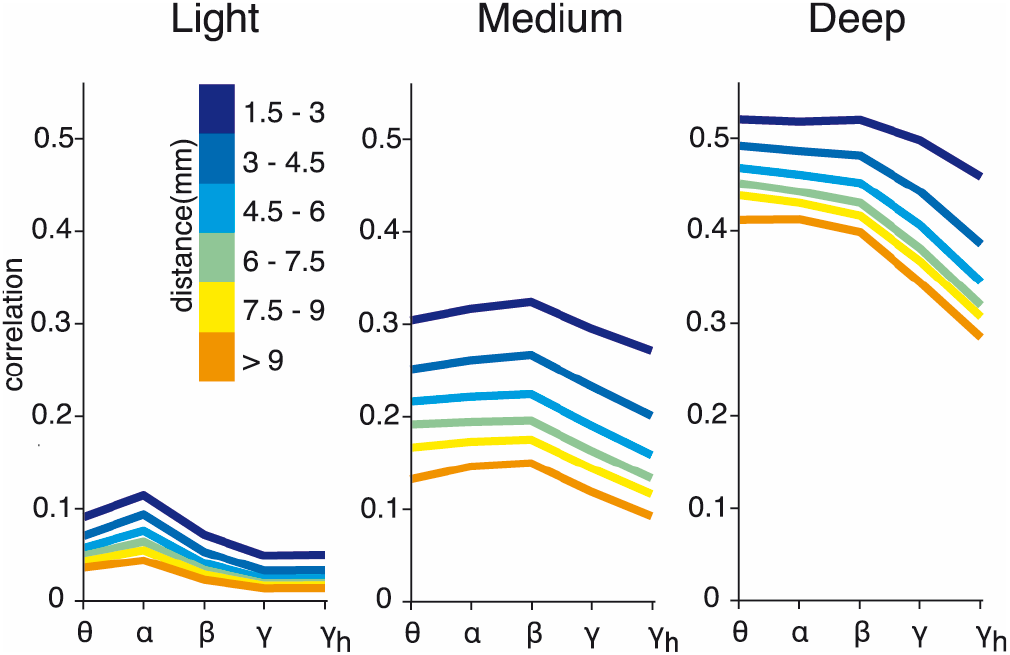
Mean amplitude correlation for the three network states as a function of frequency band and inter-contact distance. Correlation shows a drop-off as a function of distance and frequency for all states.

### Spatial structure of envelope ICMs remains robust to anesthesia depth and distance effects

As described above, the most obvious effect of anesthesia was an overall increase of envelope ICMs between and within all areas. The relative spatial pattern within the connectivity matrix, however, appeared similar within each frequency band across anesthesia-dependent network states. To further examine this aspect, we re-expressed each value within each area-level coupling matrix as a z-score relative to all correlation values of the particular coupling matrix. The resulting mean z-scored connectivity matrix (Fig. 5A), representing only the spatial structure of the envelope ICMs separated from the absolute coupling strength, remained similar across anesthesia depths. The topography of the envelope ICMs was similar for all frequency bands.

**Figure 5.**
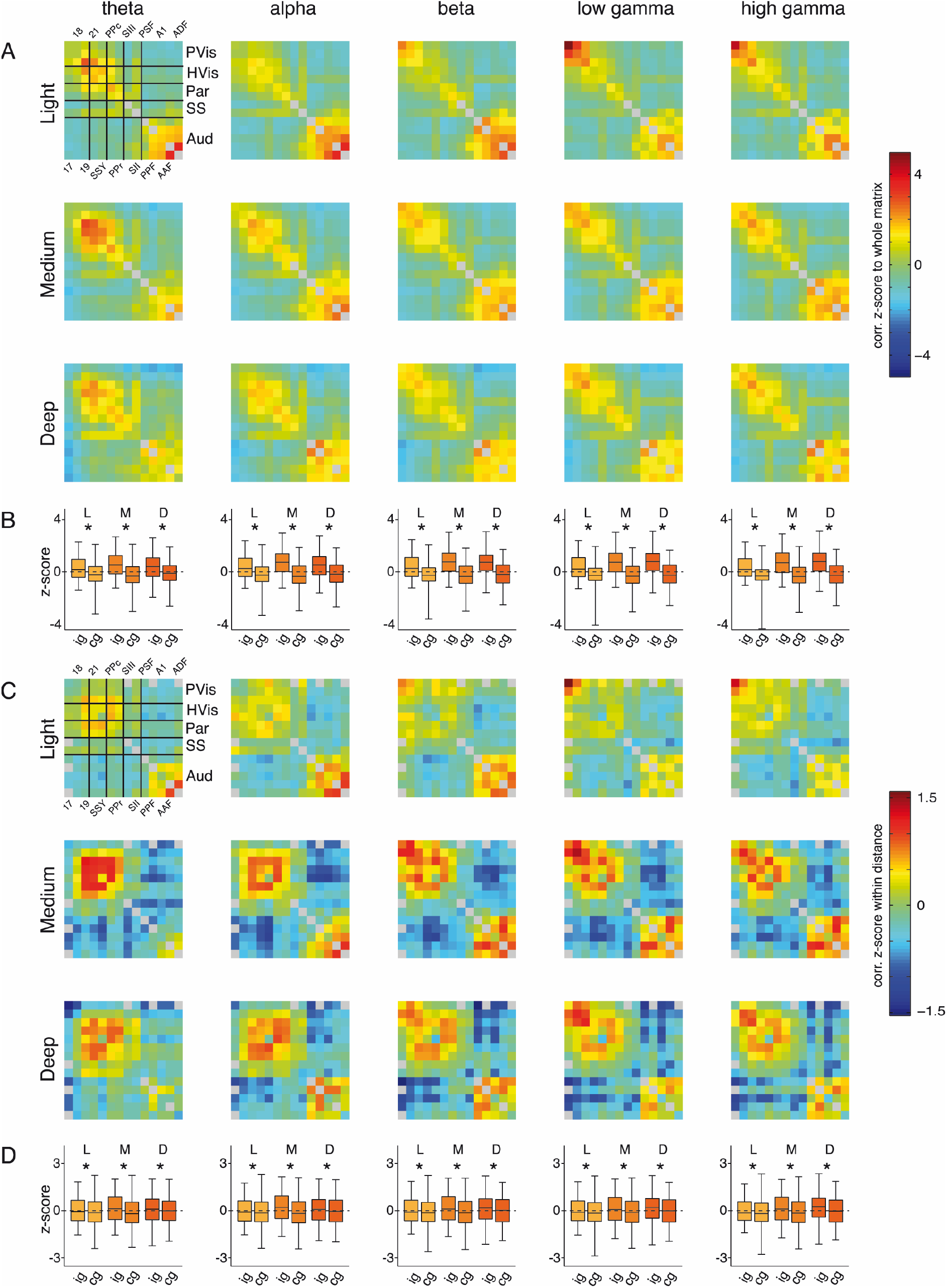
Normalized envelope ICMs show similar spatial patterns across network states. (A) Each correlation value was re-expressed as a z-score of all values in the matrix. The resulting patterns where similar across frequency bands and network states. (B) Corresponding within-group ICMs (ig) were higher compared to cross-group coupling (cg) in all three states. (C) Each correlation value was re-expressed as a z-score to the correlations of the contact pairs with a similar inter-contact distance (cf. Suppl. Fig. 3). Connectivity patterns were preserved in the resulting distance-corrected matrices. (D) Coupling within groups of cortical areas was still higher than cross-group coupling. (see next page)

Since the functional groups comprised neighbouring cortical areas, the general decrease of amplitude correlation with distance would induce a bias to higher intra- than cross-group correlations. We therefore tested whether the spatial patterns visible in the coupling matrices could be explained exclusively by the lower distance of the contacts comprised in areas and functional groups. For this purpose we re-expressed each correlation coefficient of the contact-level coupling matrix as a z-score relative to all correlations between contact pairs with the same distance (Suppl. Fig. 3). On the resulting correlation matrices, we recomputed the coupling matrices for the three network states defined above. As shown in Fig. 5C, the topographical patterns observed previously were still largely preserved. Furthermore, within-group z-scores were still higher than across-group z-scores (KS-Test, Bonferroni-Holm-corrected, p<0.05). This suggests that higher envelope ICMs between functionally related areas cannot exclusively be attributed to their spatial proximity.

### Envelope modulation frequency changes with anesthesia depth

To further investigate the temporal dynamics of amplitude envelopes in the three network states, we calculated the mean power spectrum of the amplitude envelope for each frequency band and each state (Fig. 6A). The spectrum of the signal amplitude was relatively flat for all frequency bands in the light anesthesia network state. During medium anesthesia, a peak around 0.7 Hz was visible, which was most pronounced in the gamma band. For the deep anesthesia state, the peak frequencies of the amplitude modulation were below 0.1 Hz.

**Figure 6.**
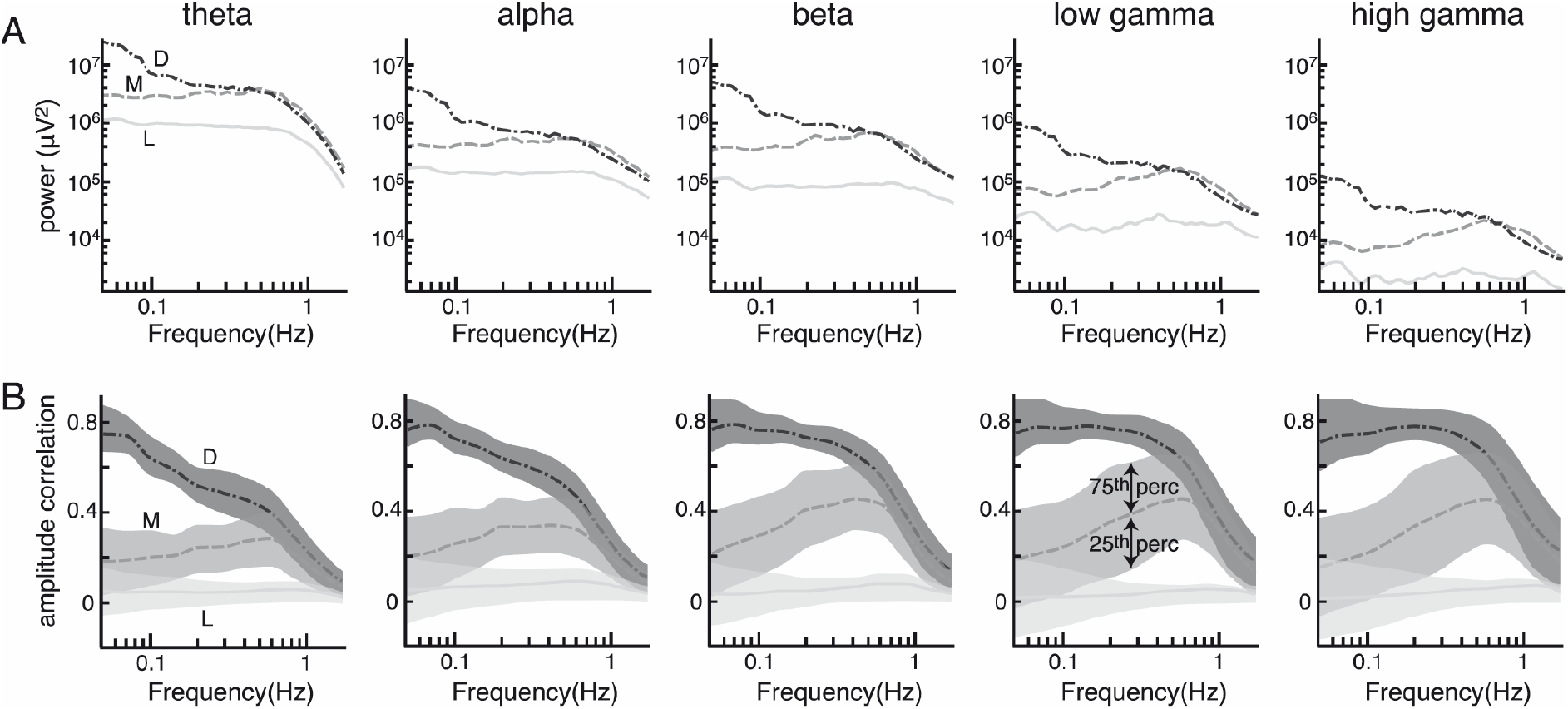
Spectral characteristics of the envelope ICMs differ across the network states. (A) Spectrum of the amplitude envelope in light (L), medium (M) and deep (D) anesthesia for the different frequency bands. In low anesthesia, the spectrum is relatively flat. In medium anesthesia a peak emerges around 0.5Hz which is most pronounced in the gamma band, and in deep anesthesia, power shifts to slower frequencies below 0.1Hz. (B) Envelope ICMs computed at different modulation frequencies of the amplitude envelope. The shaded area represents the values between the 25th and the 75th percentile. Note the correlation peak around 0.5Hz in medium anesthesia and the increased correlation at slower frequencies in deep anesthesia.

This change in modulation frequency raised the question whether amplitude correlations differed for specific modulation frequencies, and whether this dependency changed with anesthesia. We therefore bandpass-filtered the time course of the amplitude envelope in eleven frequency bands between 0.04 and 2 Hz and calculated the mean correlation within each of them for each frequency and each cluster (Fig. 6B). Correlation at all modulation frequencies increased with anesthesia. During light anesthesia, the resulting spectrum was flat. For the medium anesthesia state, it showed a peak around 0.5 Hz, which was more pronounced in the beta- and gamma- than in the theta- and alpha-band. During the deep anesthesia network state, correlation showed a constant decay with modulation frequency.

## Discussion

This study shows, using the ferret cortex as a model, that quantification of envelope ICMs allows to characterize network states which emerge across different concentrations of isoflurane anesthesia. Based on hierarchical clustering of the coupling matrices, we could differentiate three network states under light, medium and deep anesthesia. The particular anesthetic dosage at which a state was reached varied across animals, reflecting individual differences in the response to the anesthetic. Overall, our results suggest that envelope ICMs in ongoing activity reflect the functional organization of cortical networks in their topographical structure. Although the average coupling increased with anesthesia depth, the topography of functional networks was conserved across different network states. The dominant modulation frequencies of the amplitude envelopes shifted across anesthesia depths, and so did the frequency bands at which the strongest envelope ICM was observed. Anesthesia states classified by envelope ICMs were associated with different dynamics of local neural population activity visible in the electrocorticogram. The light anesthesia state with low global correlations was characterized by continuous activity patterns, the high correlation levels of the deep anesthesia state were related to low-frequency burst suppression.

### Methodological considerations

The network we examined consists of primary as well as higher auditory, visual and somatosensory areas, thus comprising a relatively large area of the cortex. However, since frontal and occipital areas show different reactions to anesthesia (Cimenser et al. 2011; Supp et al. 2011; Sellers et al. 2013), our results cannot be generalized on the entire neocortex without further experimental evidence. The approach for quantification of the envelope ICMs used here includes the orthogonalization of the phases of the filtered signals which eliminates effects of volume conduction, but comes at the price of also removing physiological in-phase activity, which plays an important role in neural processing (Singer 1999; Engel et al. 2001). As shown recently, this also implies that our approach underestimates functional connectivity (Hipp et al. 2012). Our state classification by hierarchical clustering interprets deepening anesthesia as switches between clearly distinguishable states. Due to the relatively large logical distance between the three clusters visible in the dendrograms, we believe this approach to the data is justified, even though it ignores minor within-state differences of the coupling patterns. Furthermore, our approach does not take into account spontaneous fluctuations in network connectivity strength over time (de Pasquale et al. 2010).

### Changes of local dynamics and spectral characteristics under anesthesia

The ECoG-signal can predominantly be attributed to the synchronous population activity of cortical pyramidal neurons (Buzsáki et al. 2012). We observed a change of the ECoG-signal from continuous activity under light anesthesia to burst suppression patterns under higher isoflurane concentrations, an effect long known from surface EEG recordings in clinical settings (Brown et al. 2010). Spectral characterization of local activity revealed changes known from previous studies that have employed isoflurane anesthesia (Sellers et al. 2013). With increasing isoflurane concentration we observed a relative increase in theta, alpha and beta power and a relative decrease in gamma power. These changes in the local dynamics relate to potentiation of GABAergic inhibition in the thalamo-cortical system (Lukatch et al. 2005). At the single-cell level, bursts correspond to epochs of phasic depolarization, while periods of suppression correspond to electrical silence resulting from anesthetic-mediated suppression of glutamatergic transmission (Steriade et al. 1994; Lukatch et al. 2005).

The spectrum of the amplitude envelopes changes across anesthesia depth, likely representing the drastic alteration of activity pattern visible in the surface ECoG signal. The continuous spontaneous activity of the low anesthesia state was reflected in envelopes predominantly modulated at frequencies below 0.1 Hz, similar to those described for the awake brain (Leopold et al. 2003; Nir et al. 2008). In medium anesthesia, the envelope spectrum showed an overall power increase with a peak around 0.6 Hz especially for the alpha-, beta-, and gamma-band, likely a spectral fingerprint of the emerging burst suppression-patterns. The further increase of modulation frequency especially below 0.1 Hz in deep anesthesia might be the correlate of the sparser occurrence and longer duration of bursts, however a more systematic examination of burst properties would be necessary to verify this assumption. The correlation of the envelopes is highest at the dominant modulation frequencies of the particular state.

### Envelope ICMs and cortical network state

Complementary to the states of local neuronal population activity visible in the patterns of the raw ECoG signal, we defined network states through amplitude envelope correlations. With deepening anesthesia we observed a global increase of coupling strength with constant relative spatial patterns. The spatial specificity of envelope ICMs we observed across anesthesia depths has been described before in the awake state and sleep. Homologous areas display high interhemispheric correlations in envelope ICMs calculated from MEG (Hipp et al. 2012). Within the same hemisphere, amplitude correlations between ECoG-contacts within the sensorimotor cortex have been observed to be higher than those to areas of other modalities in the awake state, slow-wave- and REM-sleep (He et al. 2008). On a finer spatial scale, the resemblance of tonotopy by envelope ICMs has been shown in the macaque (Fukushima et al. 2012). To our knowledge, our study is the first to examine high-resolution envelope ICMs at different depths of isoflurane anesthesia.

In the medium and deep anesthesia network states, which were characterized by burst suppression patterns, cortical activity was not equally coupled across the whole set of recorded areas, but amplitude correlation was still higher for neighboring cortical sites than for distant ones. Correspondingly, synchrony and similarity of bursts have been shown to depend on cortical distance. Bursts occur nearly simultaneously in adjacent cortical areas (Contreras et al. 1997). However, large-scale ECoG recordings in humans have shown that burst suppression is not completely homogeneous across the whole cortex (Lewis et al. 2013). First, not all areas enter and exit burst suppression simultaneously. Second, under burst suppression, not all bursts are global, and the global ones are less synchronized for more distant cortical areas. These results are well compatible with our observation that even under deep burst suppression, the cortex is not homogeneously synchronized, but the spatial and functional structure of the network is preserved.

In the awake brain, amplitude envelopes and their correlations have been suggested to remain stable across behavioral states (Leopold et al. 2003). Our results show that anesthesia modulates some of their basic characteristics, while others remain constant: While mean correlation increases with anesthesia depth, it shows a negative dependence on distance between recording sites in all three states. Similarly, a drop-off of envelope ICMs for all frequency bands as a function of distance has been described in the awake rhesus monkey (Leopold et al. 2003). Furthermore, we observed a drop-off of correlation as a function of frequency, which suggests more local amplitude co-modulation of higher frequencies. In contrast, in the awake brain especially high coherence values have been reported for amplitude correlations of the gamma band (Leopold et al. 2003). Whether this is due to the different scale of our recordings, which cover a larger area of much smaller ferret brain, caused by the orthogonalization, or a property of the anesthetized brain remains open to further investigation.

### Relation to fMRI studies on functional connectivity under anesthesia

To our current understanding, connectivity revealed by amplitude correlation of electrophysiological signals is comparable to that defined by BOLD correlation in restingstate fMRI (Leopold and Maier 2012; Engel et al. 2013). The low-frequency modulation of neural activity represented in the amplitude envelope is tightly linked to the temporal dynamics of local cerebral blood flow and oxygen extraction mirrored in the BOLD signal via the hemodynamic response function (Martin 2014). ECoG-amplitude- and BOLD-derived correlations of ongoing activity are highly correlated (He et al. 2008; Schölvinck et al. 2010; Keller et al. 2013), and resting networks defined by independent component analysis (ICA) of MEG power envelopes resemble those known from fMRI studies (de Pasquale et al. 2010, 2012). The close association of ECoG amplitude envelopes and cerebral blood flow via the hemodynamic response function has been described in isoflurane-anesthetized rats and is preserved in the anesthetized state (Liu et al. 2011).

However, it should also be noted that the comparability of fMRI studies to electrophysiological results is limited by some methodological differences. Most resting-fMRI studies segment activity patterns into distinct networks using spatial ICA or examine networks of few, sometimes distant seed regions. Due to the limited temporal resolution of fMRI they typically do not consider frequencies above 0.1 Hz, which we show to contribute to amplitude correlations under anesthesia. For the same reason, these studies usually do not examine the temporal dynamics of the signals.

Since different anesthetics potentially differ in their effects on brain connectivity (Brown et al. 2010; Williams et al. 2010; Jonckers et al. 2013; Grandjean et al. 2014), we confine our discussion to studies using isoflurane. It has been shown that the spatial specificity of fMRI-coupling is preserved under isoflurane anesthesia (Vincent et al. 2007; Hutchison et al. 2010). An increase of functional connectivity with deepening anesthesia was observed in a study examining BOLD patterns in the sensorimotor cortex of the rat under three isoflurane concentrations (1.0%, 1.5%, 1.8%) (Liu et al. 2013). Distinct sensorimotor subnetworks where distinguishable under the lowest isoflurane concentration and BOLD correlations decreased as a function of distance. With increasing dosage, spatial specificity of correlation patterns decreased and high brain-wide correlations emerged. The same study described a similar effect on power correlations from three recording sites, again in accordance with our own findings (Liu et al. 2013).

These strong long-range BOLD correlations observed under deep anesthesia have been shown to be associated with electrophysiological burst suppression and to break down under isoflurane concentrations leading to an isoelectric local field potential (Liu et al. 2011). High global correlations have been described under 1.5% isoflurane, which is likely comparable to our medium- or deep-anesthesia state (Kalthoff et al. 2013). While there are no previous electrophysiological data on envelope ICMs under isoflurane anesthesia, increased interhemispheric amplitude correlations have been observed during sleep (Nir et al. 2008). Although these findings are well compatible with our results, other studies report a breakdown of functional connectivity with increasing isoflurane concentration (Wang et al. 2011; Hutchison et al. 2014), and similar observations have been made under other anesthetics (Lu et al. 2007; Bettinardi et al. 2015).

Taken together, it is difficult to derive a consistent picture of resting fMRI networks under anesthesia from the available literature. These contradictory findings might be due to variations in experimental and analytical methodology. Experimental protocols differ in species, regions between which coupling is calculated, isoflurane concentrations, and comedication. This makes an assignment of similar brain states across studies very difficult. In addition, neurovascular coupling and thus the BOLD signal are altered by blood CO_2_- and O_2_-level (Nasrallah et al. 2015), a mechanism by which different ventilation protocols might influence the observed coupling patterns. Additionally, different fMRI preprocessing strategies seriously influence the observed functional connectivity. Since high correlations in deep anesthesia are based on synchronous activity across large brain areas, they disappear if the global brain signal is removed in the preprocessing of the fMRI data, giving the opposed impression of sparsely correlated networks (Kalthoff et al. 2013; Liu et al. 2013).

### Envelope ICMs and cortical network function

At first glance, it might seem counterintuitive that deepening anesthesia leads to increased functional connectivity. However, increased envelope ICMs in deep anesthesia are associated with the rather stereotypic local patterns of burst suppression, which are highly synchronized throughout the network, as already suggested by Liu (Liu et al. 2011). Both the more stereotyped local dynamics and the overall increase in functional connectivity are in line with previous hypotheses on loss of consciousness under deep anesthesia (Alkire et al. 2008; Supp et al. 2011). It has been suggested that anesthesia has two effects on the cortex: a breakdown of integration, and the entrainment to stereotypic activity patterns leading to a loss of information (Alkire et al. 2008). Our data agree with this notion and support the following interpretation. While the topographic patterns of the envelope ICMs remain stable, the signals propagating across the network change drastically with the transition into deeper anesthesia. In the awake state and under light anesthesia, dynamics of ongoing activity is complex and external stimuli elicit differentiated responses representing stimulus features. Under burst suppression, specific local responses are replaced by stereotypic bursts similar to those occurring spontaneously (Hudetz and Imas 2007). These evoked bursts spread to areas otherwise unresponsive to stimuli of the particular modality (Hudetz and Imas 2007; Land et al. 2012). Thus, on the one hand the stereotypic patterns prevailing throughout the network hinder the emergence and propagation of ‘informative’ intrinsic and stimulus-related neural activity. On the other hand, global connectivity increases, possibly leading to a reduction of the functional segregation of areas. Both aspects lead to a reduction in functional network complexity which may account for the loss of consciousness under anesthesia (Alkire et al. 2008).

Besides these considerations on anesthesia, our results reinforce the hypothesis that envelope ICMs closely reflect patterns of structural connectivity between the underlying areas (Engel et al. 2013), robust even to a severe perturbation as that induced by anesthesia. For resting-fMRI, the high similarity of functional to structural connectivity has been shown by tracer injection (Vincent et al. 2007). Nonetheless, as suggested previously, envelope ICMs may have a role in shaping processing capacities of functional networks (Engel et al. 2013). Contrary to classical views, ongoing activity is likely to carry information and to be endowed with meaningful spatiotemporal structure, which reflects previous learning and can bias the processing of stimuli (Engel et al. 2001; Deco et al. 2011). Envelope ICMs seem to represent coherent excitability fluctuations that may lead to coordinated changes in the activation of brain areas. As we have proposed, this might regulate the availability of neuronal populations or regions for participation in an upcoming task and gate the emergence of more specific interactions for processing of stimulus- or response-related signals (Engel et al. 2013). However, these aspects clearly await further investigation.

## Acknowledgements

This work was supported by grants from the Deutsche Forschungsgemeinschaft (DFG), SFB 936/A1/A2 and SPP 1665 EN533/13-1. We thank Dorrit Bystron for excellent technical support.

